# Learning Perturbation-specific Cell Representations for Prediction of Transcriptional Response across Cellular Contexts

**DOI:** 10.1101/2023.03.20.533433

**Authors:** Gal Keinan, Karen Sayal, Alon Gonen, Jiang Zhu, Lena Granovsky, Jeremy England

## Abstract

High-throughput screens (HTS) are widely utilized to profile transcriptional states across multiple cell types and perturbations, and are often the first step on the bridge to the patient. However, their representative capacity to encompass all the cellular contexts encountered in a patient is limited. Thus, we present PerturbX, a novel deep learning model that leverages the rich information obtained from HTS to predict transcriptional responses to chemical or genetic perturbations in unobserved cellular contexts, and demonstrate its effectiveness in an experimental setting. Furthermore, we show that the model is able to uncover interpretable genetic signatures associated with the predicted response, which can ultimately be translated into the clinical setting.

## 1. Introduction

Information-rich phenotypes provide a detailed picture of the cellular consequences of chemical or genetic perturbations which, in specific contexts, may have a downstream effect on cellular fitness. In particular, *gene expression* profiles provide a robust and informative phenotypic measure of cellular responses to perturbations. High-throughput screens (HTS) are utilized to profile transcriptional states across various cellular contexts under different perturbations (McFarland et al., 2020). However, their representative capacity remains limited relative to the vast combinatorial landscape of all cells and perturbation pairs. This highlights the need to utilize machine learning tools to predict the outcome of unseen experiments.

Moreover, as the availability of diverse pre-clinical and clinical datasets for precision oncology is expanding, an important unresolved question remains as to how best to integrate insights gained between the pre-clinical and clinical settings. One component will involve leveraging the information richness of high-throughput perturbational screens to identify and validate signatures which can be effectively generalized to different contexts. The ultimate goal will be to predict the phenotypic impact of treatment (i.e. perturbation) in a patient based on markers obtained from the pre-clinical realm.

To this end, we propose PerturbX, a novel deep learning model trained to predict the transcriptional effect of a given perturbation in a range of cellular contexts (e.g. different cell types), in which the perturbational response has not been experimentally observed. PerturbX is based on an encoder - decoder architecture that learns a mapping between the unperturbed state representation of cells to the transcriptional effect of a given perturbation. The unperturbed state is represented by the unperturbed gene expression profiles of cell lines, and the transcriptional effect is represented by the vector of differential expression. We show that the model effectively uncovers interpretable factors of variation within the unperturbed state which are associated with the observed patterns of transcriptional response. These “biomarkers” of response could ultimately be mapped between the pre-clinical and clinical settings, bridging the gap between the two domains.

Our main contributions are:

1. We introduce PerturbX and demonstrate its ability to successfully predict transcriptional responses to perturbations in unseen cellular contexts using data readily available in the public domain.
2. We show that PerturbX can learn biologically meaningful and interpretable representations of cell types.
3. We propose a method for identifying the predictive features, or biomarkers, of response captured by PerturbX.

## 2. PerturbX

PerturbX is a deep learning model trained to predict the transcriptional effect of a perturbation across cellular contexts by learning a *perturbation-specific* cell type representation.

This representation captures the complex transcriptional similarities between different cell types in response to a specific compound or genetic perturbation, and is inferred from features of the unperturbed state of a cell. The proposed method allows for generalization on experimentally unobserved cell types.

The input to the model consists of the population of *unperturbed* (DMSO-treated) single cells from a specific cell type along with a perturbation encoding. The target is the average differential expression (DE) of that cell type in response to the given perturbation.

More formally, denote the sets of cell types and perturbations by *C* and *P*, respectively. For each (*c, p*) ∈ *C* × *P*, let *X*^(*c,p*)^ be the subset of the training data consisting of gene expression profiles of single cells of type *c*, that have been perturbed using the perturbation *p*. Also, let *X*^(*c*)^ be the subset of *unperturbed* single cells of type *c*. Denote their respective averages by 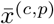 and 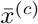.

The *input* to the model consists of the pair (*X*^(*c*)^, *e*_*p*_), where *e*_*p*_, the *perturbation encoding*, is a one-hot vector representing the perturbation *p*.

The *target* is the mean differential expression vector

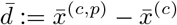

Note that our formulation falls in the category of *multiple instance learning*, where for each cell type-perturbation pair, (*c, p*), we have a set of instances (unperturbed expression of *K* single-cells) as the input, and a single target vector (namely, the DE) associated with this set. By using the entire single-cell population as input, the model is able to utilize the information about the *distribution* for the prediction of the differential expression. In the sequel we often omit specific mention of (*c, p*) if no confusion arises.

### Bootstrap

To simplify the training and induce stochasticity, on each epoch we replace *X*^(*c*, *p*)^ and *X*^(*c*)^ by two i.i.d. samples

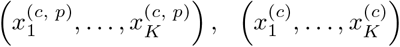

drawn uniformly from from *X*^(*c,p*)^ and *X*^(*c*)^, respectively.

#### 2.1. Architecture

The model architecture, shown in Figure 1, is based on an encoder-decoder network. However, unlike classical autoencoders, we do not reconstruct the input, but rather predict a high-dimensional response vector (i.e., the differential expression).

**Figure 1.**
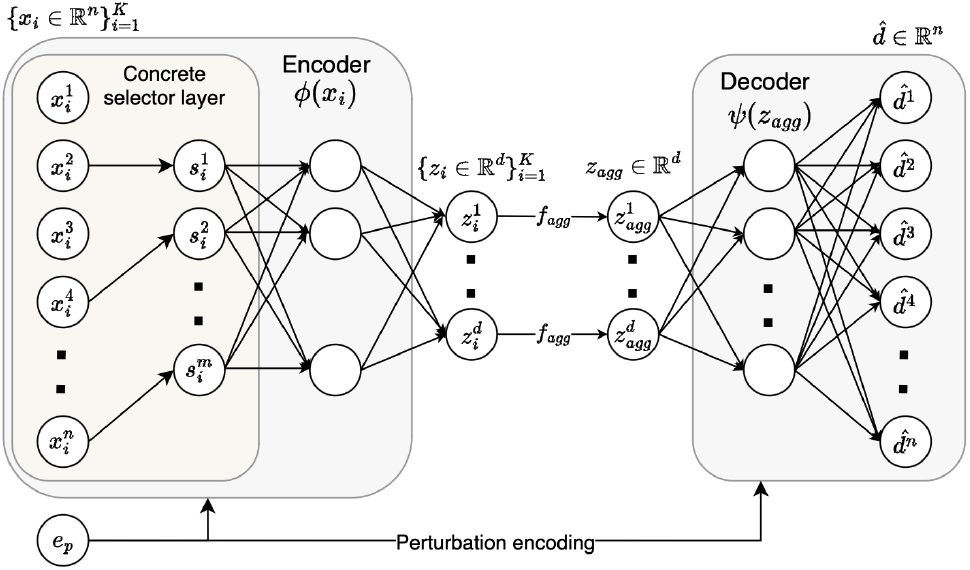
Schematic of the PerturbX model architecture. Given *K* single cells from a specific cell type and a perturbation encoding as input, the model’s encoder selects a subset of the input genes using the concrete selector layer and maps each of the cells into the latent space. The *K* different embeddings are aggregated to summarize the latent representation for that cell type. The aggregated representation is then mapped by the decoder into a prediction of the post-perturbation differential expression.

Formally, the encoder *ϕ* maps each pair (*x*_*i*_, *e*_*p*_) for *i* = 1, …, *K* into a latent vector *z*_*i*_. The latent vectors are then aggregated using 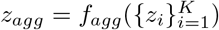 (see discussion below on the choice of the aggregation function *f*_*agg*_). Finally, the decoder *ψ* maps the pair (*z*_*agg*_, *e*_*p*_) into a prediction, 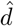, of the post-perturbation differential expression.

A key component in the encoder’s architecture is the *concrete selector* layer (Maddison et al., 2016; Balin et al., 2019) on which we elaborate in Section 2.2.1.

### Aggregation Strategies

We explore two aggregation strategies: 1) *mean* aggregation, where we simply take the average over the latent embeddings in the set, and 2) *attention*-based aggregation (Ilse et al., 2018), in which we compute a weighted average over the latent embeddings where weights are determined by a learnable function. For both of these choices, our model can be written as

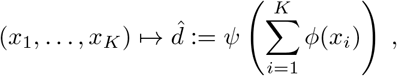

where the aggregation weights are absorbed into the encoder *ϕ*. This ensures that the model is *permutation-invariant* (Zaheer et al., 2017).

### Loss

The loss w.r.t. a single pair of input and target is

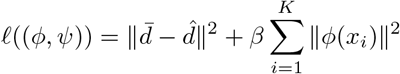

where *β* is a hyperparameter controlling the complexity of the latent representation.^1^

#### 2.2. Model Explainability

A key aspect of the perturbation response prediction is the ability to explain the predicted outcome. This can be done by identifying the factors of variation in the input which contribute the most to the predicted outcome. In our case, this corresponds to identifying the genes whose upor downregulation has a strong effect on the model’s response prediction, or equivalently, on the latent representation of the different cell types.

Our approach consists of two steps. First, we narrow down the feature space using a differentiable feature selection layer, trained end to end with the rest of the model, and second, we rank the selected features based on their importance for the response prediction.

##### 2.2.1. CONCRETE SELECTOR LAYER

We use concrete random variables to select the important input genes in a differentiable way which is jointly optimized with the rest of the model parameters for the response prediction task. Similarly to Balin et al. (2019), we use samples from a Gumbel-Softmax distribution which are smoothly annealed into a categorical distribution throughout model training. The choice of annealing schedule should allow for exploration of different gene selections in the early phases of training, and converge to an informative set of genes towards the end of training.

Whilst this approach contributes to model explainability, ablation studies also demonstrate improvement in model performance relative to a fully-connected network.

##### 2.2.2. GENE RANKING

To gain better insight on the relative importance of the selected genes (with respect to a specific perturbation *p*), we examine their effect on the aggregated latent representation *z*_*agg*_, as changes in *z*_*agg*_ reflect axes of variation along the manifold of predicted differential expression.

Denote by *Z* ∈ ℝ^|*C*|*×d*^ the matrix whose rows correspond to aggregated latent representations of the different cell types under the perturbation *p*. We decompose *Z* into its principal components, and quantify the effect of the input genes in a specific principal axis by projecting *Z* onto this axis, and estimating gene importance using Shapley values.

Ultimately, the set of genes recovered by this process can be utilized to construct a complex biomarker of response to *p*, which is transferable to other domains (e.g., complex cell models, patients).

It is important to note that by inspection of the variability along a chosen principal component, one can associate a subset of input genes with specific changes in the response.

## 3. Experiments

To demonstrate the performance of PerturbX, we used the pooled single cell RNA sequencing data generated by McFarland et al. (2020). The data consists of post-perturbation single cell gene expression profiles for 24 cell lines under various compounds, including DMSO treatment as a negative control. We excluded perturbations for which the single-cell populations were too small to reflect their underlying distribution. The expression data was normalized and *log*(1 + *x*)-transformed. Finally, we subsetted the expression profiles to 5000 highly-variable-genes.

For detailed exploration, we focus our analysis here on the small molecule inhibitor, idasanutlin, as a representative compound. This is due to the selective and heterogeneous nature of the responses it induces across different cell lines, which makes the prediction task more challenging.

### 3.1. Predicting the Response to Idasanutlin

Idasanutlin blocks the interaction of MDM2 with p53 (Ding et al., 2013). p53 is an established tumor suppressor (Baker et al., 1989). MDM2 binds to p53 resulting in the enzymatic degradation of p53 (Momand et al., 1992). Idasanutlin binds directly to MDM2, blocks the MDM2-p53 interaction and thereby restores the tumour suppressive properties of p53 (Ding et al., 2013). *TP53* is the gene encoding the p53 protein. Many cancer cell lines have inactivating mutations in *TP53* which prevent them from responding to idasanutlin (Michaelis et al., 2011). Of the 24 cell lines in our dataset, 17 cell lines are *TP53*-mutated and show a weak response to idasanutlin. In contrast, a pronounced transcriptional response is observed in the *TP53* wild-type (WT) cell lines.

Our encoder, *ϕ*, consists of a concrete selector layer that selects 64 genes, followed by two fully-connected layers to produce an 8-dimensional latent embedding. The decoder, *ψ*, consists of two fully-connected layers.

For evaluation, we split the data into 6 folds, each consists of 4 different cell lines, stratified according to *TP53* mutational status. We hold out each single fold and train the model on the remaining folds. Finally, we report the *R*^2^ and MSE between the measured and predicted post-perturbation expression profiles, averaged over the folds. Since most genes do not vary significantly in response to perturbation, we evaluate our metrics on the top 100 differentially expressed genes (DEGs). This ensures that we capture the prediction quality of the actual effect, without being masked by noise from unresponsive genes. However, for a complete evaluation of model performance, we also report the MSE on the entire gene set.

We benchmarked the predictive performance of PerturbX to 1) a baseline model that discards cell line information and predicts the average effect (DE) over all the training cell lines, and 2) scGen (Lotfollahi et al., 2019), which utilizes a conditional VAE and latent space arithmetics to predict the perturbed single cell distribution in unseen cell types.

Whilst there are more recent models designed for perturbation response prediction (e.g., Hetzel et al. (2022), which is designed to genralize to unseen compounds), we chose scGen since it is specifically aimed for generalizing to *unobserved cellular contexts*. Additionally, we note that scGen is a generative model that predicts the post-perturbation single-cell distribution whereas PerturbX is focused on the average effect. Hence, for comparison purposes, we only use the mean expression predicted by scGen.

Figure 2 shows a scatter-plot of the true vs predicted *differential expression* profiles of response to idasanutlin generated by PerturbX (left), the baseline model (center), and scGen (right) in two unobserved cell lines from one of the 6 folds - NCIH226 (*TP53* WT) and BICR6 (*TP53* mutated). It demonstrates the model’s ability to distinguish responsive and non-responsive cell lines and to make accurate predictions of the transcriptional effect. scGen, by design, averages the effect in the latent space over all observed cell lines and performs comparably with the baseline.

**Figure 2.**
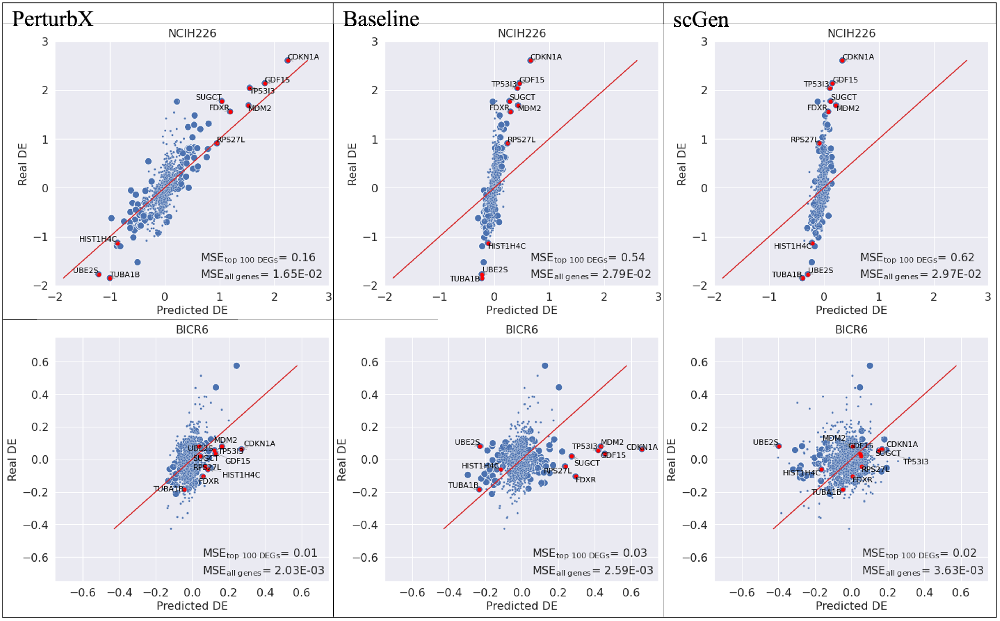
PerturbX successfully predicts *differential* gene expression in cell lines which do (NCIH226) and do not (BICR6) respond to idasanutlin. The top 100 DEGs (indicated by large dots) were chosen based on the response across all cell lines. The data points highlighted in red indicate the top 10 DEGs.

Averaged *R*^2^ and MSE over the folds are shown in Table 1. Noticeably, the improvement in prediction accuracy obtained by PerturbX over the other models is more significant in the *TP53*-WT cell lines which are more responsive, and consequently, harder to predict. Furthermore, as the baseline model predicts the mean DE, taken uniformly over the training cell lines, the improvement over this baseline is attributed to our model’s ability to exploit cell line *similarity*.

**Table 1.**
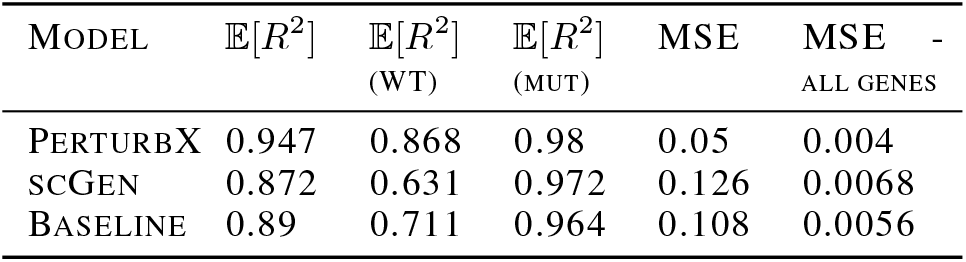
Performance metrics for PerturbX, scGen and the baseline model. *R*^2^ results are reported on all cell lines, and separately for *TP53*-WT and *TP53*-mutated cell lines. All metrics are measured over the top 100 DEGs unless stated otherwise.

To investigate how PerturbX can capture a biologically representative latent embedding, we next examined the 2D UMAP of the trained latent representation *Z*. The UMAP plot, shown in Figure 3a, confirms that cell lines, in both train and test sets, are mapped into different clusters in the latent space in accordance with their *TP53* mutational status.

**Figure 3.**
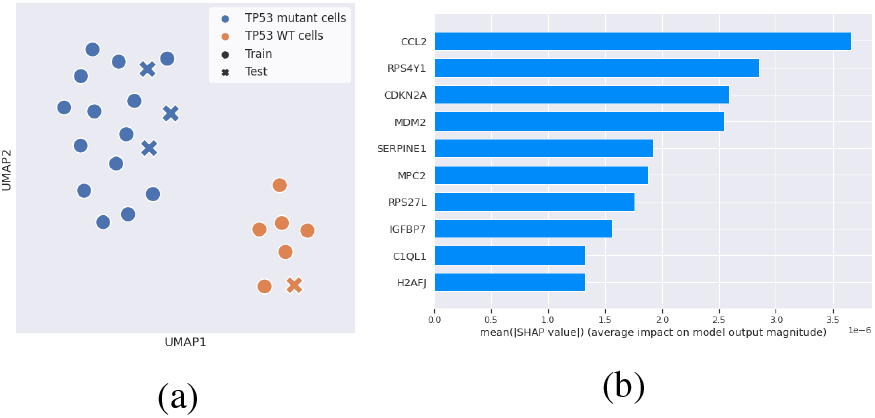
(a) UMAP visualization of the trained embedding *Z*. Each dot is the aggregated representation of a single cell line. (b) Genes contributing to the predicted expression upon perturbation of p53 signaling are enriched for known p53 targets.

We next determined the contribution of the input genes on the predicted outcome (see section 2.2.2). Decomposition of *Z* using PCA reveals that the first principal component, *w*_1_, captures the distinction between responsive and nonresponsive cell lines. Therefore, to identify the genes which are indicative of cell response, we compute gene importance scores with respect to *w*_1_ by assessing the Shapley values of the function *X* → *ϕ*(*X*) · *w*_1_. The top 10 genes are shown in Figure 3b. 6 of these genes are known *TP53* targets: *CCL2* (Tang et al., 2012), *MDM2* (Momand et al., 1992), *SERPINE1* (Akula et al., 2020), *RPS27L* (He & Sun, 2007), *IGFBP7* (Chen et al., 2011), and *C1QL1* (Mei et al., 2008).

## 4. Discussion

We present PerturbX, a deep learning model which predicts the transcriptional response to perturbation based on the unperturbed state representation of a given cellular context. It learns a low-dimensional representation of cell types that reflects the similarity in response to a given perturbation, and is able to capture a diverse and heterogeneous response landscape. In addition, PerturbX identifies genetic features from the unperturbed state which most significantly contribute to the predicted output in a manner aligned with a priori domain-specific knowledge.

## Footnotes

1 In our experiments, rather then using the standard Euclidean norm, we used the norm 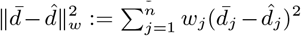, where 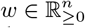 is a fixed weight vector assigning larger weights to more dominant genes.

